# Divergent Therapeutic and Prognostic Impacts of Immunogenic Features in Undifferentiated Soft Tissue Sarcoma and Myxofibrosarcoma

**DOI:** 10.1101/2025.03.27.645727

**Authors:** Siddh van Oost, Debora M. Meijer, Zeynep B. Erdem, Marieke E. IJsselsteijn, Jessica Roelands, Suk Wai Lam, Melissa S. Boejharat, Brendy E.W.M. van den Akker, Ruud van der Breggen, Inge H. Briaire-de Bruijn, Lukas J.A.C. Hawinkels, Anouk A. Kruiswijk, Manon van der Ploeg, Pauline M. Wijers-Koster, Rick L. Haas, Michiel A.J. van den Sande, Noel F.C.C. de Miranda, Judith V.M.G. Bovee

## Abstract

Myxofibrosarcoma and undifferentiated soft tissue sarcoma (USTS) are genetically complex soft tissue sarcomas with distinct morphological features. Treatment typically involves surgery, often combined with neoadjuvant chemo- or radiotherapy. To better understand the immunobiology of these sarcomas and its associations with treatment response and prognosis, we performed transcriptomic and immunophenotypic profiling.

RNA sequencing was performed on 13 USTS and 10 myxofibrosarcomas and immunological profiles were compared with soft tissue sarcoma data from The Cancer Genome Atlas (*n* = 206 including 44 USTS and 17 myxofibrosarcomas). Immune contextures were further evaluated in 16 USTS and 11 myxofibrosarcomas using imaging mass cytometry. Characterization of T cell and macrophage infiltration in tumors was further assessed in 23 USTS and 22 myxofibrosarcomas through multispectral immunofluorescence and immunohistochemical analysis.

USTS and myxofibrosarcomas demonstrated immunogenic features compared to other soft tissue sarcomas, with subsets of USTS and myxofibrosarcomas demonstrating high T cell infiltration while USTS demonstrated a higher infiltration by myeloid cells as compared to myxofibrosarcoma. Prognostically, T cells and CD68^+^CD163^+^ macrophages were associated with metastasis-free survival in USTS but not in myxofibrosarcomas. Notably, in USTS, neoadjuvant radiotherapy appeared to induce cytotoxic T cell infiltration and depletion of myeloid cells, whereas these effects were not observed in myxofibrosarcomas.

These findings highlight important differences in the immunobiology of USTS and myxofibrosarcomas with therapeutic and prognostic implications. These differences should be taken into account given the growing availability of immunotherapeutic options for treating patients with soft tissue sarcomas.

## Introduction

Soft tissue sarcomas (STS) are a rare and diverse group of malignant mesenchymal neoplasms that predominantly arise in the extremities or trunk. Undifferentiated soft tissue sarcoma (USTS) and myxofibrosarcoma are two subtypes distinguished by their morphological features and lack of specific line of differentiation. USTS, previously classified as undifferentiated pleomorphic sarcoma (UPS) and undifferentiated spindle cell sarcoma, present exclusively as intermediate- or high-grade deep-seated tumors, while myxofibrosarcomas encompass a spectrum of low- to high-grade tumors and can occur either superficially or deep-seated. Localized disease is typically treated with surgery, often complemented by (neo-)adjuvant radiotherapy depending on tumor size, grade, and anatomical location (1). However, clinical outcomes remain poor, with approximately 10% of patients experiencing local relapse and up to 50% distant metastases (2). As options for systemic therapy in recurrent disease are limited, improving clinical management through the development of novel therapeutic approaches is crucial.

Genomically, USTS and myxofibrosarcoma present complex and heterogeneous profiles characterized by extensive chromosomal alterations and few recurrent mutation targets (e.g., *TP53, ATRX, RB1*) (3). Additionally, these tumors display similar transcriptional and epigenetic profiles when compared to other soft tissue sarcomas (4, 5). Despite these overlaps, their clinical behavior differs considerably: USTS is more prone to metastasize, with distinct responses to neoadjuvant therapy, whereas myxofibrosarcomas show a higher propensity for local recurrence, suggesting underlying biological differences (2, 6). The contribution of the immune microenvironment to these differing outcomes remains unclear (7).

To address this, we characterized the immune microenvironment of USTS and myxofibrosarcomas using transcriptomic and immunophenotypic profiling. Immune-related transcriptional profiles were compared with STS from The Cancer Genome Atlas (TCGA) and immune cell populations were associated with metastasis-free and disease-specific survival. Moreover, the effect of radiotherapy on the immune microenvironment was explored. T cell and CD68^+^CD163^+^ macrophage infiltration correlated with metastasis-free survival in USTS, but not in myxofibrosarcoma. Notably, radiotherapy in USTS increased cytotoxic T cell phenotypes while reducing myeloid populations, a response not observed in myxofibrosarcomas. These findings underscore distinct immunobiological features between USTS and myxofibrosarcomas, with potential implications for developing tailored therapeutic strategies.

## Methods

### Patient samples

Formalin-fixed paraffin embedded (FFPE) and snap-frozen material was collected for 16 USTS and 18 myxofibrosarcoma patients diagnosed between 2008 and 2021. This cohort comprised high- and low-grade myxofibrosarcomas, with patients receiving surgery alone, post-operative radiotherapy or pre-operative radiotherapy. Subsequently, the cohort was expanded to include FFPE biopsy samples from an additional 14 USTS and 15 myxofibrosarcoma patients diagnosed between 2011 and 2023. These cases were exclusively high-grade tumors that underwent neoadjuvant radiotherapy. All samples in this study were pseudo anonymized and handled according to the ethical guidelines outlined in ‘Code for Proper Secondary Use of Human Tissue in The Netherlands’ by the Dutch Federation of Medical Scientific Societies. A waiver of consent was obtained from the medical ethical evaluation committee (Medisch-Ethische Toetsingscommissie Leiden Den Haag Delft; protocol number: B17.036 and B20.067). As a result, this study adheres to the Declaration of Helsinki.

For imaging mass cytometry, regions representative of the tumor’s immune microenvironment were identified on Haematoxylin & Eosin (H&E)-stained sections of FFPE tissue by a soft tissue tumor pathologist (JVMGB). Tissue Microarrays (TMAs) were then constructed using a TMA Master (3DHISTECH). Each resection contributed four cores (1.6 mm in diameter), with two central and two peripheral cores, while biopsies provided two cores. For immunohistochemistry/immunofluorescence, FFPE blocks of biopsies with sufficient tumor tissue were selected for whole slide imaging. Pathological response to radiotherapy was assessed by two soft tissue tumor pathologists (JVMGB & SWL), who defined the modified EORTC response score including percentage of necrosis, hyalinization/fibrosis and viable tumor adding up to 100% of tumor volume (8, 9).

### RNA sequencing

Sample processing and sequencing of RNA was performed as described previously on treatment-naïve tumors of 13 and 10 USTS and myxofibrosarcoma respectively (10). In short, RNA was extracted from snap-frozen tissue using TRIzol and isopropanol/ethanol, followed by purification with the RNeasy kit, including DNase treatment. Paired-end 150 base pair (bp) reads were generated on a NovaSeq6000 Illumina at GenomeScan (Leiden, The Netherlands). Sequencing data were processed with the RNA-seq BioWDL pipeline from the SASC (BioWDL Github, LUMC, The Netherlands). Reads were aligned to the hg38 reference genome using STAR and gene expression was quantified using HTSeq-count (11). The processed data were stored as a matrix containing gene counts per sample.

### Integration with TCGA-STS dataset

RNA sequencing data from 206 revised cased of primary STS from the TCGA (3) were downloaded and processed using TCGA Assembler (R, v.2.0.3). This dataset included USTS (*n* = 44), myxofibrosarcoma (*n* = 17; including 3 low-grade), dedifferentiated liposarcomas (*n* = 50), soft tissue leiomyosarcomas (*n* = 53; including 11 low-grade), uterine leiomyosarcomas (*n* = 27; including 1 low-grade), malignant peripheral nerve sheath tumors (*n* = 5) and synovial sarcomas (*n* = 10). Gene symbols were converted to HGNC gene symbols and Entrez Gene identifiers using biomaRt (R, v.2.60). The dataset was filtered for overlapping genes with our own RNA sequencing dataset, referred to as the Leiden Center for Computational Oncology (LCCO) dataset, using Entrez Gene identifiers. Since the TCGA data was in transcripts per million (TPM), the processed RNA sequencing counts from the LCCO cohort were converted to TPM by using IOBR (R, v.0.99.8), which also removed genes without HGNC symbols. The LCCO and TCGA datasets were then merged based on HGNC symbols. Normalization followed the method described by Roelands *et al*. (12). It was performed within lanes and between lanes by using EDASeq (R, v.2.38), after which the data was quantile normalized with preprocessCore (v.1.66.0) and log2 transformed.

### Gene expression analysis

Batch correction was performed with limma (R, v.3.60.0) to compare immune-related gene expression between our dataset and the TCGA dataset. As described previously, the immunologic constant of rejection (ICR) gene signature was utilized to characterize the Th1-like inflammatory state of the STS microenvironment (13, 14). Additionally, the STS were evaluated with MCPcounter (R, v.1.2.0), which clusters samples into previously established Sarcoma Immune Classes (SIC) by using the TCGA-STS probe set (15). Z-Scores were calculated per gene/cell population to visualize the immune-related transcriptional signatures using ComplexHeatmap (R, v.2.20.0). The attributed clusters were visualized per subtype, per dataset in bar plots with ggplot2 (R, v.3.5.1).

### Imaging mass cytometry

The conjugation of BSA-free antibodies and immunodetection using a 40-marker tumor immune microenvironment panel (**Supplementary table 1**) were carried out as previously described (10, 16). Briefly, four-µm sections of the generated TMAs were deparaffinized, rehydrated and subjected to heat-induced antigen retrieval in 1x low pH antigen retrieval solution (pH 6, Thermo Fisher Scientific). The first set of primary antibodies was incubated overnight at 4°C. Following PBS washes supplemented with 1% BSA and 0.05% Tween, sections were incubated with conjugated anti-mouse (Abcam, ab6708) and anti-rabbit (Abcam, ab6701) antibodies for one hour at room temperature. Next, the second batch of primary antibodies was applied for five hours at room temperature, followed by an overnight incubation at 4°C with the final batch. Finally, DNA was stained using 1.25 µM Iridium DNA intercalator (Standard BioTools) for five minutes. ROIs of 1000×1000 µm were ablated with a Hyperion mass cytometry imaging system (Standard BioTools) at the Flow Core Facility (LUMC, Leiden). Imaging data was acquired with CyTOF Software (v.7.0) and exported using MCD Viewer (v.1.0.5).

**Table 1.** Clinicopathological characteristics of the studied cohorts of USTS and myxofibrosarcomas. Tumor grades were assessed according to the FNCLCC grading system. The immuno?uorescence (IF) and immunohistochemistry (IHC) cohort included 7 myxofibrosarcomas (MFS) and 9 USTS from the whole transcriptome sequencing (RNAseq) and imaging mass cytometry (IMC). IQR = interquartile range, RT = radiotherapy.

| Variable | RNAseq & IMC |  | IF & IHC |  |
| --- | --- | --- | --- | --- |
|  | MFS | USTS | MFS | USTS |
| **Total patients, *n*** | 18 | 16 | 22 | 23 |
| **Gender, *n* (%)** |  |  |  |  |
| Female | 6 (33.3) | 8 (50) | 9 (40.9) | 10 (43.5) |
| Male | 12 (66.7) | 8 (50) | 13 (59.1) | 13 (56.5) |
| **Age (years), median (range)** | 64 (24-77) | 74 (40-80) | 70 (24-81) | 67 (40-87) |
| **Follow-up (months), median (range)** | 52 (6-150) | 50 (12-92) | 24 (4-63) | 41 (15-104) |
| **Grade, *n* (%)** |  |  |  |  |
| Low (1) | 5 (27.8) | - | - | - |
| High (2, 3) | 13 (72.2) | 16 (100) | 22 (100) | 23 (100) |
| **Anatomical location, *n* (%)** |  |  |  |  |
| Head & Neck | 1 (5.56) | - | - | 1 (4.3) |
| Upper extremities | 4 (22.2) | 1 (6.3) | 3 (13.6) | 4 (17.3) |
| Trunk | 4 (22.2) | 1 (6.3) | 3 (13.6) | - |
| Lower extremities | 9 (50) | 14 (87.5) | 16 (72.7) | 18 (78.3) |
| **Max tumor size (cm), median (IQR)<sup>a</sup>** | 6.95 (7.65) | 8.9 (3.88) | 10 (7.45) | 8.9 (7.45) |
| **Treatment, *n* (%)<sup>b</sup>** |  |  |  |  |
| Adjuvant radiotherapy | 2 (11.1) | - | - | - |
| Neoadjuvant radiotherapy<sup>c</sup> | 10 (55.6) | 16 (100) | 22 (100) | 23 (100) |
| Surgery alone<sup>d</sup> | 6 (33.3) | - | - | - |
| **Histological response to neoadjuvant RT (%), median (IQR)** |  |  |  |  |
| Vital tumor | 60 (10) | 15 (34.3) | 49.5 (45) | 10 (30.5) |
| Hyalinization / fibrosis | 20 (15) | 40 (43.3) | 15 (15) | 55 (40) |
| Necrosis | 12.5 (20) | 20 (37.5) | 25 (54.6) | 20 (25) |
| **Pathological response to neoadjuvant RT, *n* (%)<sup>e</sup>** |  |  |  |  |
| Good | - | 6 (37.5) | 5 (22.7) | 9 (39.1) |
| Poor | 6 (100) | 10 (62.5) | 17 (77.3) | 14 (60.9) |
| **Tumor depth, *n* (%)** |  |  |  |  |
| Deep | 9 (50) | 16 (100) | 14 (63.6) | 23 (100) |
| Superficial | 9 (50) | - | 8 (36.64) | - |
| **Patients with recurrent disease, *n* (%)** |  |  |  |  |
| Local recurrence | 3 (16.7) | 2 (12.5) | 2 (9.1) | 1 (4.3) |
| Metastasis | 4 (22.2) | 8 (50) | 10 (45.5) | 13 (56.5) |

a: size was missing for one USTS from the face
b: treatment for primary tumors
c: one high-grade MFS patient received neoadjuvant RT + pazopanib
d: one high-grade MFS patient had an amputation
e: pathologic response was defined as <5% vital tumor after neoadjuvant radiotherapy

### Imaging mass cytometry analysis

Analysis of the imaging mass cytometry data was performed as described previously (10, 16). In short, images were normalized in Matlab (v.R2021a) and binarized in Ilastik (v.1.3.3). Probability masks were generated in Ilastik and used for cell segmentation in CellProfiler (v.2.2.0). Single cell-FCS files were generated in ImaCyte (17) and utilized for phenotyping in CytoSplore (v.2.3.1). Mean cell density per mm^2^ was calculated for every sample. Lineage markers used for phenotyping are shown in **Supplementary table 2**. The relative abundance of phenotypes was calculated using the total number of cells per sample and visualized using ComplexHeatmap.

### Multispectral immunofluorescent imaging

A T cell Opal panel comprising four markers was developed for immunofluorescent imaging of pre-treatment biopsies using a Vectra system (Akoya Biosciences) as described previously (10, 18). The panel included PD-1 (D4W2J, 1:250, Cell Signaling Technology, #86163), CD8α (D8A8Y, 1:500, Cell Signaling Technology, #85336), CD3ε (EP449E, 1:1000, Abcam, ab52959) and CD4 (EPR6855, 1:2000, Abcam, ab133616). Each antibody was paired with a specific Opal fluorophore in an optimized staining sequence to maximize intensity and specificity: CD8-Opal 690 (1:100), CD4-Opal 570 (1:400), PD-1-Opal 620 (1:100) and CD3-Opal 520 (1:400). Endogenous peroxidase was blocked with 0.3% H_2_O_2_ in methanol, antigen retrieval was performed with citrate buffer (pH 6.0) and tissue was blocked with SuperBlock™ Blocking Buffer (ThermoFisher Scientific). Slides were incubated with primary antibodies for one hour at room temperature, washed with PBS-Tween (0.05%,) and incubated with BrightVision DPVO-HRP (Immunologic) for 10 minutes. After washing, the first Opal reagent (Akoya Bioscience), diluted in Opal amplification diluent, was applied for 10 minutes. Slides were then heated in citrate buffer (pH 6.0) for 15 minutes at a reduced wattage and washed with PBS-Tween. This process was repeated for the remaining antibodies. Finally, slides were incubated with DAPI (1:1000) for five minutes and mounted with ProLong™ Gold Antifade Mountant (ThermoFisher Scientific). Up to five ROIs per slide were selected based on consecutive H&E slides and imaged at 20x magnification. Image processing was performed using inForm (v.2.4) and analyzed with QuPath (v.0.3.1). Due to nonspecific staining, CD4 was excluded from the analysis. T cells were classified as CD8^+^ or CD8^-^, and further categorized based on PD-1 expression (PD-1^+^ or PD-1^-^). Cell counts per image (1.5 mm x 2 mm) were normalized per patient as counts per mm^2^ tissue. Patients were stratified into T cell high and T cell low clusters based on the median total T cell count per subtype.

### Double immunohistochemical staining

Double immunohistochemical (IHC) staining was performed to identify macrophages in pre-treatment biopsies using the ImmPRESS Duet Double Staining Polymer Kit (HRP Anti-Mouse IgG-brown, AP Anti-Rabbit IgG-magenta, Vector Laboratories, MP-7724-15). The markers CD163 (10D6, Mouse, 1:400, MONOSAN, MONX10445) and CD68 (D4B9C, Rabbit, 1:3200, Cell Signaling Technology, #76437) were applied on the same slide following the manufacturer’s protocol. Slides were scanned using a Pannoramic™ 480 (3D HISTECH) and analyzed with QuPath. Macrophages were segmented by using optimal sum density for cell detection, excluding most CD68^+^ and/or CD163^+^ tumor cells. The same ROIs were selected as for the T cell imaging, and macrophage counts per ROI were normalized to counts per mm^2^ tissue. Patients were classified in the same manner as for T cell counts, based on CD68^+^ and CD68^+^CD163^+^ phenotypes.

### Survival analysis

Kaplan-Meier survival curves were generated using the survminer (v.0.4.9) and survival (v3.6-4) packages in R. Disease-specific survival was defined as the time until death due to the disease. Given the high number of censored cases in the cohort, statistical significance was assessed using the log-rank *P* value. Univariate and multivariate Cox proportional hazards analyses were conducted in R, with only variables found significant in the univariate analysis included in the multivariate model.

### Data availability

The RNA sequencing data generated in this study are publicly available in Gene Expression Omnibus GEO at GSE285944. The IMC data is available in BioStudies at S-BIAD1555. The code used to process data is published on the LUMC Bone & Soft Tissue Pathology Group GitLab (https://git.lumc.nl/bstp/papers/immunogenic-features-of-usts-and-myxofibrosarcoma). All other data are available from the corresponding author upon reasonable request.

## Results

### Clinicopathological characteristics are prognostic in myxofibrosarcoma, but not in USTS

In this study, 63 patients were examined, comprising 30 cases of USTS and 33 cases of myxofibrosarcoma. Clinicopathological data are summarized in **Table 1**. Clinicopathological investigations related to prognosis were limited to 54 patients with high-grade tumors who underwent standard-of-care treatment, including neoadjuvant radiotherapy. In USTS, no clinical parameters, including pathologic response after neoadjuvant treatment (<5% vital tumor), were found to associate with metastasis-free or disease-specific survival (**Supplementary table 3**). In contrast, tumor size and tumor depth were identified as prognostic factors for both metastasis-free and disease-specific survival in myxofibrosarcoma (**Supplementary table 4**). Interestingly, a good pathological response to radiotherapy in myxofibrosarcoma was negatively associated with metastasis-free and disease-specific survival.

Pathological responses to radiotherapy differed between the two sarcoma types. While USTS primarily exhibited hyalinization and fibrosis, myxofibrosarcoma predominantly displayed necrosis (**Supplementary figure 1A**). Furthermore, histological responses in myxofibrosarcoma were observed exclusively in large, deep-seated tumors, suggesting an association between tumor depth, tumor size, and necrosis (**Supplementary figure 1B**). Multivariate analysis did not identify any of these clinical parameters as independent prognostic factors in myxofibrosarcoma (**Supplementary table 4**).

### Heterogeneity in immune-enriched microenvironments of USTS and myxofibrosarcoma

To evaluate the “natural” immune microenvironment of USTS and myxofibrosarcoma, we analyzed treatment-naïve samples, including pre-treatment biopsies or surgical resections from patients who had not received neoadjuvant radiotherapy. A total of 23 samples (19 biopsies and 5 resections) of 13 USTS and 10 myxofibrosarcoma patients were analyzed using whole transcriptome sequencing, while 29 samples (21 biopsies and 8 resections) of 14 USTS and 15 myxofibrosarcomas were examined using imaging mass cytometry. Among these, 20 samples (15 biopsies and 5 resections) of 11 USTS and 9 myxofibrosarcomas were included in both analyses. Transcriptional profiles were compared to publicly available data for 206 primary STS from the TCGA (3, 19). Immune-related transcriptional signatures were analyzed using the ICR signature (14) (**Figure 1A**) and the SIC classification system (15) (**Supplementary figure 2A**).

**Figure 1.**
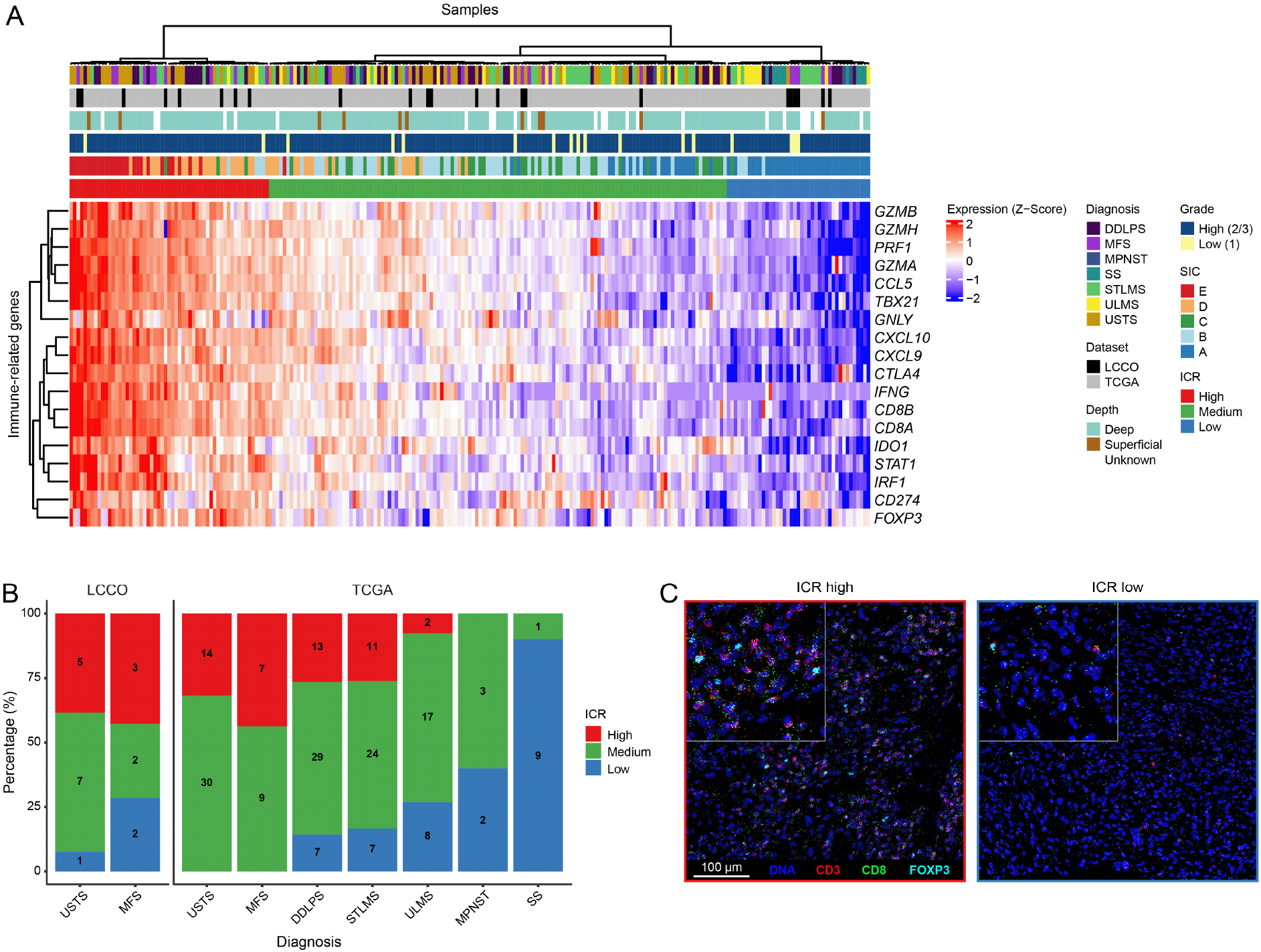
Overview of the ICR signature in USTS and myxofibrosarcoma. A) Heatmap presenting the gene expression Z-scores of 18 ICR-genes in our samples and The Cancer Genome Atlas (TCGA) samples. Annotations include diagnosis, dataset, tumor depth, tumor grade, Sarcoma Immune Class (SIC) and ICR cluster classifications. B) Bar plots illustrating the percentage of the ICR classification for each diagnosis across dataset, excluding low-grade tumors. The number of samples in each cluster is indicated within the bars. C) Representative imaging mass cytometry images of an ICR-high and an ICR-low USTS. Abbreviations: DDLPS = dedifferentiated liposarcoma; MFS = myxofibrosarcoma; MPNST = malignant peripheral nerve sheath tumor; SS = synovial sarcoma; STLMS = soft tissue leiomyosarcoma; ULMS = uterine leiomyosarcoma; LCCO = Leiden Center for Computational Oncology; ICR = immunologic constant of rejection.

Based on the ICR signature, USTS (*n* = 57, including 44 from the TCGA) and high-grade myxofibrosarcomas (*n* = 23, including 16 from the TCGA) frequently exhibited a Th1-related immune signature compared to other STS. Up to 33% of USTS and 43% of high-grade myxofibrosarcomas were classified as ICR-high, compared to only 0–27% in other STS (**Figure 1B**). Moreover, only 2% of USTS and 9% of high-grade myxofibrosarcomas presented as ICR-low, compared to as much as 14-90% in other STS (**Figure 1B**). The SIC classification further corroborated these findings, showing an enrichment of SIC D and E classifications (immune-high) in USTS and high-grade myxofibrosarcomas, alongside DDLPS in the TCGA data. These classifications were much more frequent in these tumors compared to other STS (44% in USTS, 43% in high-grade myxofibrosarcoma, 43% in DDLPS while only 0-26% in other STS; **Supplementary figure 2B**).

Myxofibrosarcomas included more tumors classified as highly vascularized (SIC C) compared to USTS (26% vs 14%; **Supplementary figure 2B+C**). However, both subtypes exhibited some heterogeneity. Up to 42% of USTS and 30% of myxofibrosarcomas were classified as SIC A (immune-desert) or SIC B (immune-low), emphasizing that, while these tumors are relatively immune-enriched, variability exists within these subtypes (**Figure 1B+C**). In contrast, low-grade myxofibrosarcomas (*n* = 3), which exhibit lower genomic complexity, displayed an immune “cold” profile overall, similar to other STS like synovial sarcoma. All three cases were classified as immune desert (SIC A) and ICR-low (**Figure 1A**; **Supplementary figure 2A**). These findings support the hypothesis that immunogenicity in sarcomas correlates with increasing genomic complexity (20, 21).

Immunophenotyping of USTS and myxofibrosarcoma by imaging mass cytometry resulted in the identification of 32 distinct cell populations, including overlapping tumor and stromal clusters due to the absence of tumor-specific markers in these sarcomas (**Supplementary table 2**). The immune cell infiltrate predominantly consisted of myeloid cells, such as macrophages and monocytes, alongside lymphoid cells, including various T cell and innate lymphoid cell populations (**Supplementary figure 3A**). Since USTS and myxofibrosarcoma tumor cells are known to express myeloid markers like CD68 and CD163, we reviewed the imaging data to ensure that the observed myeloid cell counts were not confounded by tumor cells expressing these markers. Consistent with the immune-related transcriptional profiles, low-grade myxofibrosarcomas exhibited the lowest immune cell densities (**Supplementary figure 3A+B**). Among high-grade tumors, USTS demonstrated higher immune cell densities compared to myxofibrosarcomas (**Supplementary figure 3A**), which may reflect the lower overall cellularity in myxofibrosarcomas due to their myxoid extracellular matrix.

To correct for this difference in extracellular matrix, we calculated the relative immune cell abundance per mm^2^ for all treatment-naïve samples (**Figure 2**). Unsupervised clustering revealed a relative enrichment of T cells in a subset of both myxofibrosarcomas and USTS, while myeloid cells, including monocytes and macrophages, were more prevalent in USTS compared to myxofibrosarcomas (**Figure 2**; **Supplementary figure 3C**). Tumors classified as ICR-high or ICR-medium displayed mixed levels of T cell infiltration, whereas ICR-low tumors consistently exhibited the lowest T cell infiltration. These observations align with previous findings on the relationship between T cell infiltration and the ICR score (10).

**Figure 2.**
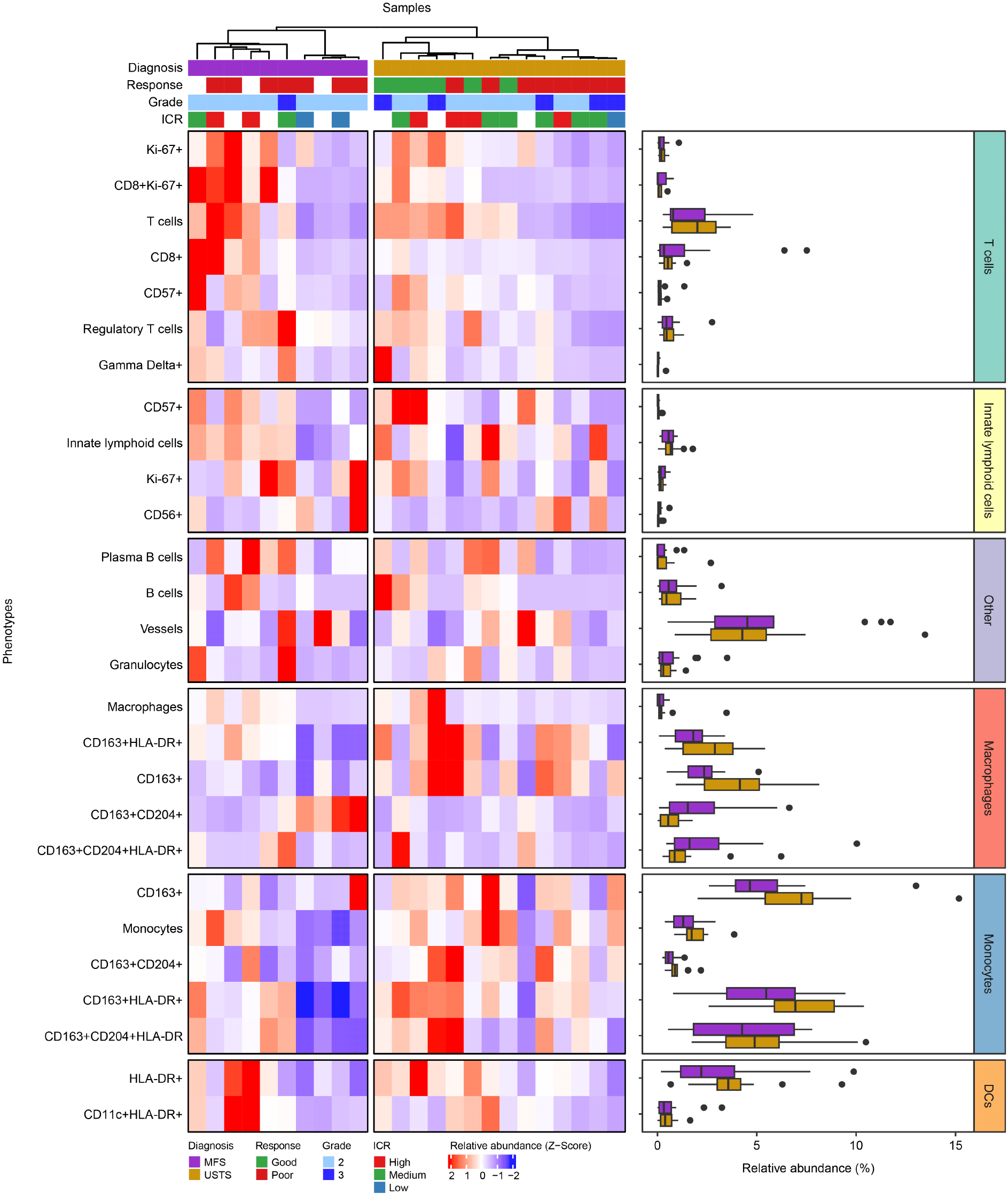
IMC-based immunophenotypic analysis of USTS and high-grade myxofibrosarcoma. The heatmap displays the relative abundance Z-scores of all identified immune cell populations within the tumor microenvironments. Samples are annotated with diagnosis, pathologic response to radiotherapy, tumor grade, and the ICR classification. Accompanying boxplots display the relative abundance of each phenotype by tumor type. Abbreviations: MFS = myxofibrosarcoma; USTS = undifferentiated soft tissue sarcoma; ICR = immunologic constant of rejection; DCs = dendritic cells.

### T cell and CD68^+^CD163^+^ macrophage infiltration is associated with an improved prognosis in USTS but not in myxofibrosarcoma

To investigate the clinical impact of T and myeloid cell populations, we composed a cohort including 45 patients that underwent the current standard-of-care treatment of neoadjuvant radiotherapy, which included 16 patients (9 USTS and 7 myxofibrosarcoma) with imaging mass cytometry data. T cell infiltration was assessed using a multispectral immunofluorescence panel, including CD3, CD8 and PD-1, while macrophages were evaluated with CD68 and CD163 two-color immunohistochemistry.

Consistent with the imaging mass cytometry results, T cell infiltration showed high variability in both USTS and myxofibrosarcoma (**Figure 3A**). A strong correlation was observed between T cell infiltration levels determined by imaging mass cytometry and those detected using multispectral immunofluorescence (**Supplementary Figure 4A**). A significantly larger proportion of CD8^+^ T cells exhibited PD-1 positivity (min: 0%, median: ~17%, max: ~87%) compared to CD8^-^ T cells (supposedly CD4^+^ T cells; min: 0%, median: ~2%, max: ~49%; *P* = 4.1e-6), with similar levels in USTS and myxofibrosarcomas (**Figure 3A**). The cohort was dichotomized by the median total T cell count within each subtype (**Figure 3B**). Higher T cell infiltration was associated with improved metastasis-free survival in USTS but not in myxofibrosarcoma (**Figure 3C**). The same association was observed for disease-specific survival, but was not found statistically significant due to the lower number of events in this category (**Supplementary figure 4B**).

**Figure 3.**
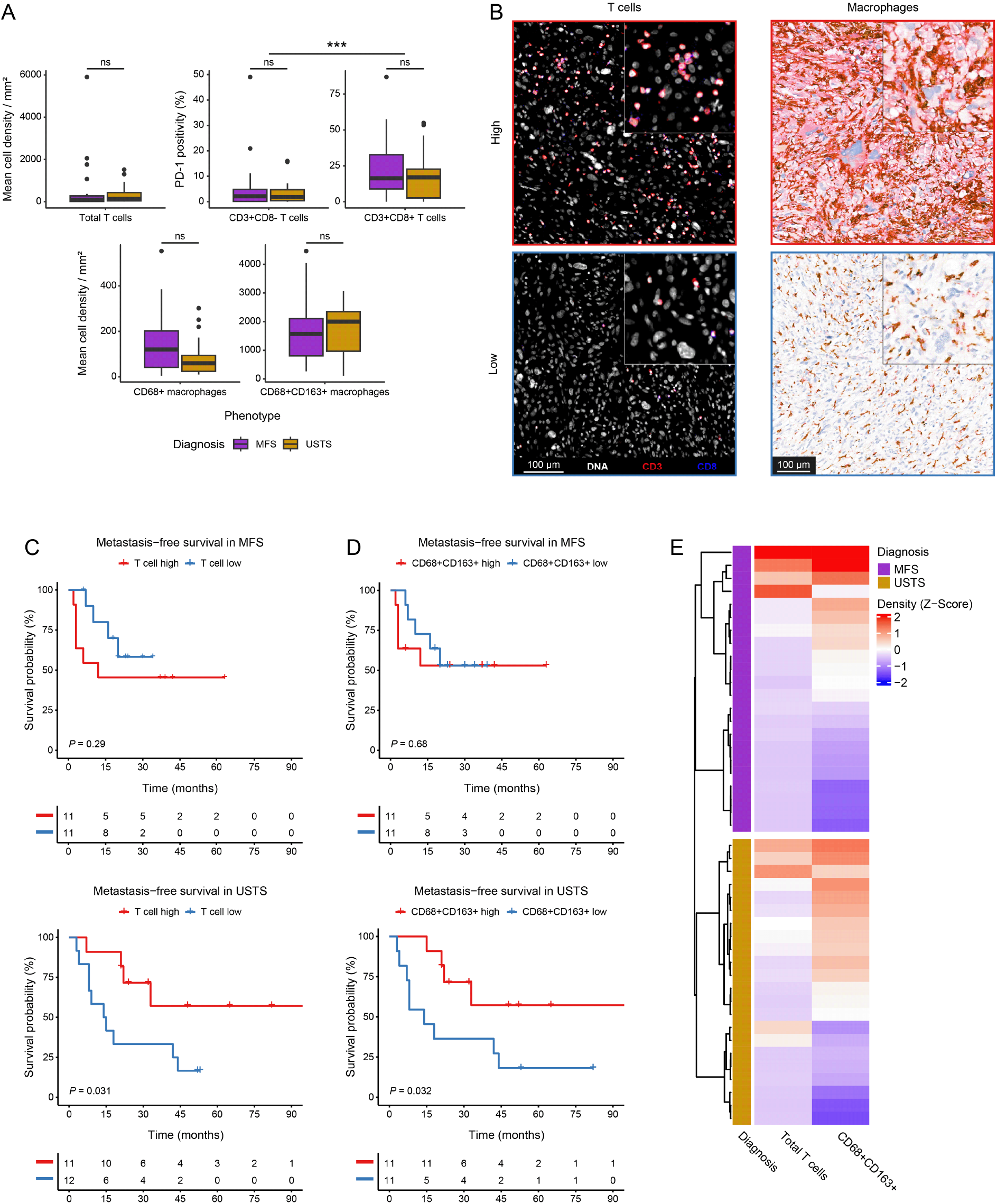
T cells & CD68^+^CD163^+^ macrophages are prognostic in USTS, but not in myxofibrosarcomas. A) Boxplots presenting the mean cell densities of T cells and macrophages, as well as the percentage of PD-1^+^ T cells (CD8^-^ and CD8^+^) in the expanded cohort. B) Example immuno?uorescence images of a T cell high and a T cell low undifferentiated soft tissue sarcoma (USTS) and immunohistochemical images of a CD68^+^CD163^+^ high and a CD68^+^CD163^+^ low myxofibrosarcoma. C) Survival curves for the metastasis-free survival in both myxofibrosarcoma and USTS separately, grouped based on T cell infiltration and D) grouped based on CD68^+^CD163^+^ macrophage infiltration. Significance is presented per Kaplan-Meier curve by the log-rank *P* value. E) Heatmap showing the association between T cell and CD68^+^CD163^+^ macrophage infiltration in both USTS and myxofibrosarcoma. The samples are annotated for their diagnosis. Abbreviations: MFS = myxofibrosarcoma.

Both subtypes exhibited comparable densities of CD68^+^CD163^+^ macrophages, while myxofibrosarcoma displayed higher levels of CD68^+^CD163^-^ macrophages, though this difference was not statistically significant (**Figure 3A**). CD68^+^CD163^+^ macrophages associated with better metastasis-free survival in USTS but not in myxofibrosarcoma (**Figure 3D**). Again, the same association was observed for disease-specific survival and not found statistically significant due to the lower number of events in this category (**Supplementary figure 4B**). Interestingly, high T cell infiltration was linked to high CD68^+^CD163^+^ macrophage infiltration in both subtypes (**Figure 3E**), suggesting coordinated roles for these immune cells in these tumors. This was further strengthened by the observation that patients with high infiltration of both T cell and CD68^+^CD163^+^ macrophages had the longest metastasis-free survival whereas patients with low infiltration of both immune phenotypes had the shortest metastasis-free survival in USTS (**Supplementary figure 4C**) Other macrophage populations were not associated with survival in either subtype (**Supplementary figure 4D**).

### Neoadjuvant radiation affects USTS and myxofibrosarcoma differently

Since T cells and macrophages are prognostic in USTS patients who received neoadjuvant radiotherapy, but not in myxofibrosarcoma patients, we studied the effect of this treatment on these immune cell populations in both subtypes. Imaging mass cytometry data was generated for matched pre- and post-treatment samples of 21 patients (13 USTS and 8 myxofibrosarcoma). In post-treatment samples, only vital tumor areas were assessed, selecting both central and peripheral regions to account for intratumoral heterogeneity. In USTS (*n* = 13), various myeloid cell populations decreased following radiotherapy (**Figure 4A**). Among T cells, CD57^+^ T cells and CD8^+^ T cells increased after radiotherapy (*P* < 0.01 and *P* < 0.05, respectively). Additional phenotypes affected by the radiotherapy in USTS included tumor/stromal cell populations and vessels, which also decreased (**Supplementary figure 5A**). These alterations were all independent from pathological response. In contrast, no significant alterations in the immune microenvironment were observed in myxofibrosarcoma following neoadjuvant radiotherapy (*n* = 8; **Supplementary figure 5A+B**).

**Figure 4.**
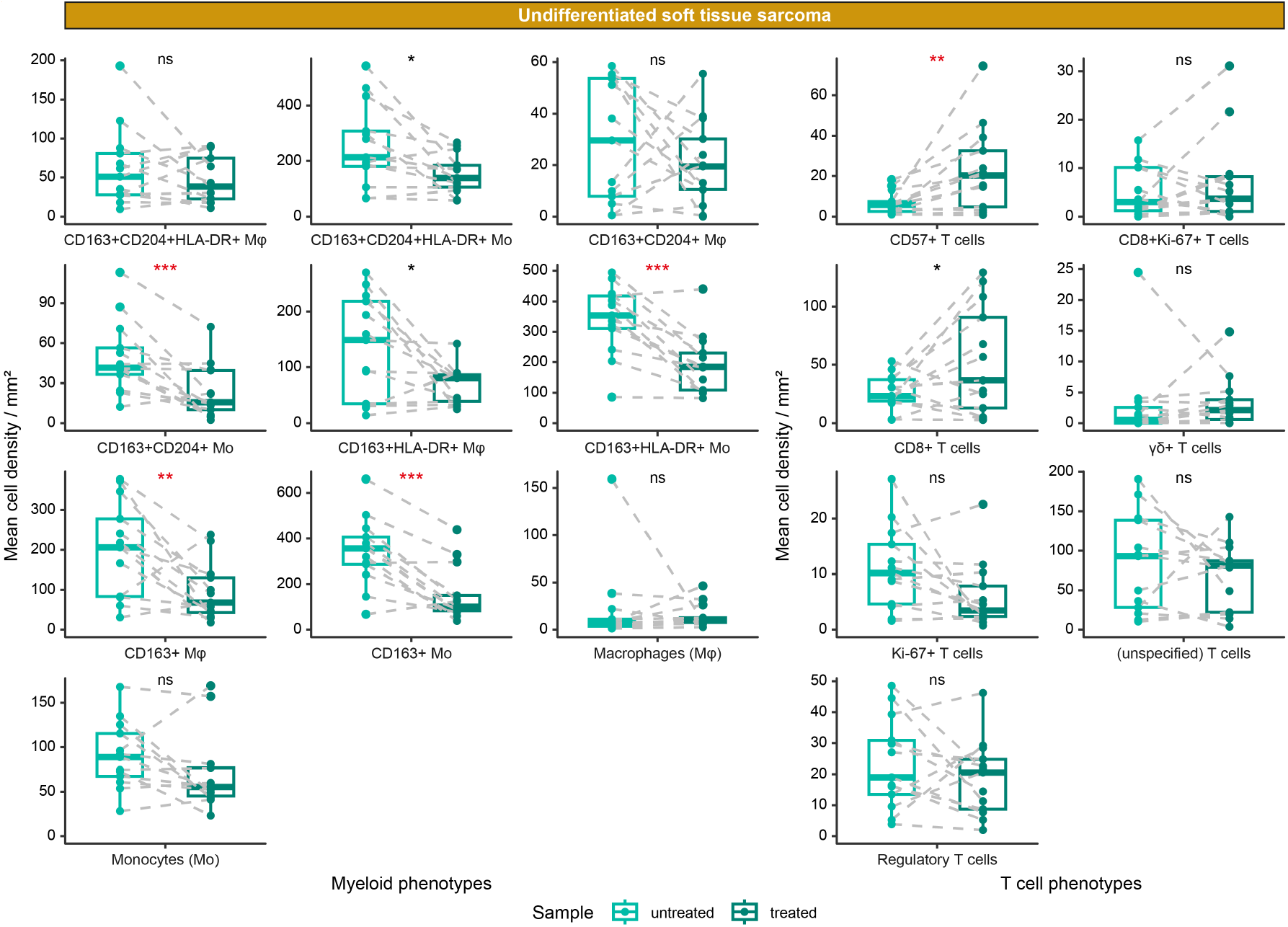
Radiotherapy affects USTS differently than myxofibrosarcoma. A) Comparison of pre-vs post-treatment undifferentiated soft tissue sarcomas (USTS), presented in paired boxplots. The significance level was evaluated with a student’s t-test followed by a Benjamini-Hochberg false discovery rate (FDR) correction. The significance is indicated per phenotype and FDR-significant phenotypes are indicated in red. ns = not significant, * = *P* < 0.05, ** = *P* < 0.01, *** = *P* < 0.001. Abbreviations: Mf = macrophages; Mo = Monocytes.

## Discussion

Although both USTS and myxofibrosarcoma demonstrate more hallmarks of anti-tumor immunity than other soft tissue sarcomas, this study reveals considerable differences between the two subtypes. While both subtypes include T cell-enriched tumors, USTS demonstrated relatively higher myeloid cell Infiltration compared to myxofibrosarcoma. In USTS, T cells and CD68^+^CD163^+^ macrophages were positively associated with improved metastasis-free survival. Interestingly, radiotherapy altered the USTS microenvironment, reducing myeloid cell populations and increasing cytotoxic T cell phenotypes. In contrast, in myxofibrosarcoma, neither T cells nor macrophages were associated with survival, and the immune microenvironment remained largely unchanged following radiotherapy. These findings highlight fundamental biological differences between the two subtypes, with USTS being more responsive to immune modulation by radiotherapy.

There is a growing interest in the immune microenvironment of soft tissue sarcomas, particularly USTS, due to its responsiveness to T cell checkpoint blockade (22, 23). However, the interaction between immunity and radiotherapy remains poorly understood. While pathological response (<5% viable tumor) was not prognostic in USTS, consistent with the findings from Danieli *et al*. (24), a pre-existing immune response appears to reduce the metastatic potential of tumors. Furthermore, we show that radiotherapy may promote a more favorable context for anti-tumor immunity in USTS, potentially enhancing T cell-mediated responses and suppressing pro-tumorigenic myeloid cell interactions. These observations appear to align with findings by Keung *et al*. (25), who observed a trend for increased T cell infiltration following radiotherapy in USTS. Interestingly, our findings contrast with the ones by Goff *et al*., who reported an increase in myeloid cells after radiotherapy in USTS (26). This discrepancy may arise from differences in data presentation; Goff *et al*. reported cell percentages relative to total cells, whereas we analyzed actual cell densities per tissue area to enable accurate pre- and post-treatment comparisons. Despite these differences, our results underscore the potential of radiotherapy to enhance T cell-mediated responses while suppressing pro-tumorigenic myeloid interactions in USTS. Although the current study highlights the prognostic effect of immunity in the context of neoadjuvant radiotherapy, it is important to note that this effect may be independent of radiotherapy altogether, as studies by Toulmonde *et al*. and Guegan *et al*. found CD8^+^ T cells to be prognostic in both untreated and chemotherapy-treated USTS respectively (27, 28).

Similar to USTS, myxofibrosarcoma has shown responsiveness to T cell checkpoint blockade (29), suggesting that anti-tumor immune responses can also be mounted in these tumors. However, our findings do not support a strong modulatory effect of radiotherapy in the immune microenvironment of myxofibrosarcomas, and pre-existing immunity appears to lack prognostic significance. This aligns with the findings of Yamashita *et al*. (21), who also demonstrated that T cell infiltration is not prognostic in myxofibrosarcoma. The divergence between the roles of the immune microenvironment in USTS and myxofibrosarcoma is striking. Despite similar overall immune cell densities, functional differences or distinct tumor-immune interactions likely drive these outcomes. For instance, Dancsok *et al*. identified SIRPα^+^ macrophages as negative prognostic markers specific to myxofibrosarcomas and not in USTS (30). Furthermore, the lack of significant immune alterations in myxofibrosarcoma post-radiotherapy underscores these differences. Future research using multidimensional approaches could provide further insights into such differences.

Neoadjuvant radiotherapy together with surgical resection effectively controlled disease in both subtypes, with only 4 out of 54 high-grade tumors experiencing local recurrence. However, metastasis remains a major challenge, occurring in approximately 50% of patients across both subtypes. This highlights the need for additional systemic therapies to target circulating or metastasizing tumor cells. Promisingly, Roland *et al*. and Mowery *et al*. demonstrated survival benefits with the combination of neoadjuvant checkpoint blockade and radiotherapy in USTS (7, 31). For cases lacking a pre-existing immune response, converting the microenvironment from “cold” to “hot” might be necessary. Guegan *et al*. observed that immune “cold” USTS responded well to neoadjuvant chemotherapy (28), which subsequently enhanced immune cell infiltration.

In conclusion, this study underscores key differences in the immune microenvironments of USTS and myxofibrosarcoma. While both subtypes exhibited high immune cell infiltration, only USTS showed improved metastasis-free survival associated with T cells and CD68^+^CD163^+^ macrophages. Radiotherapy appeared to reshape the immune response in USTS, enhancing cytotoxic T cells and reducing myeloid cell populations, yet myxofibrosarcoma remained largely resistant to such modulation. These results suggest that subtype-specific approaches may be required to improve outcomes from cancer immunotherapy. Together, these insights underscore the importance of tailored strategies to improve outcomes for these aggressive sarcomas.

## Author contributions

**S.v.O**. contributed to the conceptualization, data curation, software, formal analysis, validation, investigation, visualization, methodology, writing-original draft, project administration and writing-review editing. **D.M.M**. contributed to the conceptualization, investigation and writing-review editing. **Z.B.E**. contributed to the investigation and writing-review editing. **M.E.IJ**. contributed to the methodology and writing-review editing. **J.R**. contributed to the methodology and writing-review editing. **S.L**. contributed to the investigation and writing-review editing. **M.S.B**. contributed to the investigation and writing-review editing. **B.v.d.A**. contributed to the investigation and writing-review editing. **R.v.d.B**. contributed to the investigation and writing-review editing. **I.H.B.d.B**. contributed to the investigation and writing-review editing. **L.J.A.C.H**. contributed to the funding acquisition and writing-review editing. **A.A.K**. contributed to the resources and writing-review editing. **M.v.d.P**. contributed to the investigation and writing-review editing. **P.M.W.K**. contributed to the investigation and writing-review editing. **R.L.H**. contributed to the investigation and writing-review editing. **M.A.J.v.d.S**. contributed to the resources and writing-review editing. **N.F.C.C.d.M**. contributed to the conceptualization, supervision, funding acquisition, investigation, methodology, writing-original draft and writing-review editing. **J.V.M.G.B**. contributed to the conceptualization, supervision, funding acquisition, investigation, methodology, writing-original draft and writing-review editing. All authors have read and approved the final version of the manuscript.

## Acknowledgements

The authors would like to thank the Flow Core Facility and the Sequencing Analysis Support Core of the Leiden University Medical Center for their service.

## Funding

This work was financially supported by an unrestricted research grant from Tracon Pharmaceuticals and the intramural Leiden Center for Computational Oncology strategic fund. N.F.C.C.d.M. is funded by the European Research Council (ERC) under the European Union’s Horizon 2020 Research and Innovation Program (grant agreement no. 852832) and by the VIDI ZonMW (project number: 09150172110092).

## Disclosure of interest

The authors declare no conflicts of interest.

## Notes

### Competing Interest Statement

The authors have declared no competing interest.

### Summary of Updates

Only author affiliations have been updated.

## References

1. Gronchi A, Miah AB, Dei Tos AP, Abecassis N, Bajpai J, Bauer S, et al. Soft tissue and visceral sarcomas: ESMO-EURACAN-GENTURIS Clinical Practice Guidelines for diagnosis, treatment and follow-up. Ann Oncol. 2021;32(11):1348–65.

2. Yoshimoto M, Yamada Y, Ishihara S, Kohashi K, Toda Y, Ito Y, et al. Comparative Study of Myxofibrosarcoma With Undifferentiated Pleomorphic Sarcoma: Histopathologic and Clinicopathologic Review. Am J Surg Pathol. 2020;44(1):87–97.

3. Cancer Genome Atlas Research Network. Comprehensive and Integrated Genomic Characterization of Adult Soft Tissue Sarcomas. Cell. 2017;171(4):950–65 e28.

4. Lyskjaer I, De Noon S, Tirabosco R, Rocha AM, Lindsay D, Amary F, et al. DNA methylation-based profiling of bone and soft tissue tumours: a validation study of the ‘DKFZ Sarcoma Classifier’. J Pathol Clin Res. 2021;7(4):350–60.

5. Koelsche C, Schrimpf D, Stichel D, Sill M, Sahm F, Reuss DE, et al. Sarcoma classification by DNA methylation profiling. Nat Commun. 2021;12(1):498.

6. Imanishi J, Slavin J, Pianta M, Jackett L, Ngan SY, Tanaka T, et al. Tail of Superficial Myxofibrosarcoma and Undifferentiated Pleomorphic Sarcoma After Preoperative Radiotherapy. Anticancer Res. 2016;36(5):2339–44.

7. Mowery YM, Ballman KV, Hong AM, Schuetze SM, Wagner AJ, Monga V, et al. Safety and efficacy of pembrolizumab, radiation therapy, and surgery versus radiation therapy and surgery for stage III soft tissue sarcoma of the extremity (SU2C-SARC032): an open-label, randomised clinical trial. Lancet. 2024;404(10467):2053–64.

8. Messiou C, Bonvalot S, Gronchi A, Vanel D, Meyer M, Robinson P, et al. Evaluation of response after pre-operative radiotherapy in soft tissue sarcomas; the European Organisation for Research and Treatment of Cancer-Soft Tissue and Bone Sarcoma Group (EORTC-STBSG) and Imaging Group recommendations for radiological examination and reporting with an emphasis on magnetic resonance imaging. Eur J Cancer. 2016;56:37–44.

9. Schaefer IM, Hornick JL, Barysauskas CM, Raut CP, Patel SA, Royce TJ, et al. Histologic Appearance After Preoperative Radiation Therapy for Soft Tissue Sarcoma: Assessment of the European Organization for Research and Treatment of Cancer-Soft Tissue and Bone Sarcoma Group Response Score. Int J Radiat Oncol Biol Phys. 2017;98(2):375–83.

10. van Oost S, Meijer DM, Ijsselsteijn ME, Roelands JP, van den Akker B, van der Breggen R, et al. Multimodal profiling of chordoma immunity reveals distinct immune contextures. J Immunother Cancer. 2024;12(1).

11. Anders S, Pyl PT, Huber W. HTSeq--a Python framework to work with high-throughput sequencing data. Bioinformatics. 2015;31(2):166–9.

12. Roelands J, Hendrickx W, Zoppoli G, Mall R, Saad M, Halliwill K, et al. Oncogenic states dictate the prognostic and predictive connotations of intratumoral immune response. J Immunother Cancer. 2020;8(1).

13. Bertucci F, Finetti P, Simeone I, Hendrickx W, Wang E, Marincola FM, et al. The immunologic constant of rejection classification refines the prognostic value of conventional prognostic signatures in breast cancer. Br J Cancer. 2018;119(11):1383–91.

14. Bertucci F, Niziers V, de Nonneville A, Finetti P, Mescam L, Mir O, et al. Immunologic constant of rejection signature is prognostic in soft-tissue sarcoma and refines the CINSARC signature. J Immunother Cancer. 2022;10(1).

15. Petitprez F, de Reynies A, Keung EZ, Chen TW, Sun CM, Calderaro J, et al. B cells are associated with survival and immunotherapy response in sarcoma. Nature. 2020;577(7791):556–60.

16. Ijsselsteijn ME, van der Breggen R, Sarasqueta AF, Koning F, de Miranda NFCC. A 40-Marker Panel for High Dimensional Characterization of Cancer Immune Microenvironments by Imaging Mass Cytometry. Front Immunol. 2019;10.

17. Somarakis A, Van Unen V, Koning F, Lelieveldt B, Hollt T. ImaCytE: Visual Exploration of Cellular Micro-Environments for Imaging Mass Cytometry Data. IEEE Trans Vis Comput Graph. 2021;27(1):98–110.

18. Ijsselsteijn ME, Brouwer TP, Abdulrahman Z, Reidy E, Ramalheiro A, Heeren AM, et al. Cancer immunophenotyping by seven-colour multispectral imaging without tyramide signal amplification. J Pathol Clin Res. 2019;5(1):3–11.

19. van IJzendoorn DGP, Szuhai K, Briaire-de Bruijn IH, Kostine M, Kuijjer ML, Bovee JVMG. Machine learning analysis of gene expression data reveals novel diagnostic and prognostic biomarkers and identifies therapeutic targets for soft tissue sarcomas. PLoS Comput Biol. 2019;15(2):e1006826.

20. van Oost S, Meijer DM, Kuijjer ML, Bovee J, de Miranda N. Linking Immunity with Genomics in Sarcomas: Is Genomic Complexity an Immunogenic Trigger? Biomedicines. 2021;9(8).

21. Yamashita A, Suehara Y, Hayashi T, Takagi T, Kubota D, Sasa K, et al. Molecular and clinicopathological analysis revealed an immuno-checkpoint inhibitor as a potential therapeutic target in a subset of high-grade myxofibrosarcoma. Virchows Arch. 2022;481(4):1–17.

22. Tawbi HA, Burgess M, Bolejack V, Van Tine BA, Schuetze SM, Hu J, et al. Pembrolizumab in advanced soft-tissue sarcoma and bone sarcoma (SARC028): a multicentre, two-cohort, single-arm, open-label, phase 2 trial. Lancet Oncol. 2017;18(11):1493–501.

23. Italiano A, Bessede A, Pulido M, Bompas E, Piperno-Neumann S, Chevreau C, et al. Pembrolizumab in soft-tissue sarcomas with tertiary lymphoid structures: a phase 2 PEMBROSARC trial cohort. Nat Med. 2022;28(6):1199–206.

24. Danieli M, Barretta F, Radaelli S, Fiore M, Sangalli C, Barisella M, et al. Pathological and radiological response following neoadjuvant treatments in primary localized resectable myxofibrosarcoma and undifferentiated pleomorphic sarcoma of the extremities and trunk wall. Cancer. 2023;129(21):3417–29.

25. Keung EZ, Tsai JW, Ali AM, Cormier JN, Bishop AJ, Guadagnolo BA, et al. Analysis of the immune infiltrate in undifferentiated pleomorphic sarcoma of the extremity and trunk in response to radiotherapy: Rationale for combination neoadjuvant immune checkpoint inhibition and radiotherapy. Oncoimmunology. 2018;7(2):e1385689.

26. Goff PH, Riolobos L, LaFleur BJ, Spraker MB, Seo YD, Smythe KS, et al. Neoadjuvant Therapy Induces a Potent Immune Response to Sarcoma, Dominated by Myeloid and B Cells. Clin Cancer Res. 2022;28(8):1701–11.

27. Toulmonde M, Lucchesi C, Verbeke S, Crombe A, Adam J, Geneste D, et al. High throughput profiling of undifferentiated pleomorphic sarcomas identifies two main subgroups with distinct immune profile, clinical outcome and sensitivity to targeted therapies. EBioMedicine. 2020;62:103131.

28. Guegan JP, El Ghazzi N, Vibert J, Rey C, Vanhersecke L, Coindre JM, et al. Predictive value of tumor microenvironment on pathologic response to neoadjuvant chemotherapy in patients with undifferentiated pleomorphic sarcomas. J Hematol Oncol. 2024;17(1):100.

29. D’Angelo SP, Mahoney MR, Van Tine BA, Atkins J, Milhem MM, Jahagirdar BN, et al. Nivolumab with or without ipilimumab treatment for metastatic sarcoma (Alliance A091401): two open-label, non-comparative, randomised, phase 2 trials. Lancet Oncol. 2018;19(3):416–26.

30. Dancsok AR, Gao D, Lee AF, Steigen SE, Blay JY, Thomas DM, et al. Tumor-associated macrophages and macrophage-related immune checkpoint expression in sarcomas. Oncoimmunology. 2020;9(1):1747340.

31. Roland CL, Nassif Haddad EF, Keung EZ, Wang WL, Lazar AJ, Lin H, et al. A randomized, non-comparative phase 2 study of neoadjuvant immune-checkpoint blockade in retroperitoneal dedifferentiated liposarcoma and extremity/truncal undifferentiated pleomorphic sarcoma. Nat Cancer. 2024.

