## Supplementary materials for "Divergent Therapeutic and Prognostic Impacts of Immunogenic Features in Undifferentiated Soft Tissue Sarcoma and Myxofibrosarcoma"

**Index of supplemental information**

**Supplementary figure 1** | Association between tumor depth, tumor size and necrosis in myxofibrosarcoma

**Supplementary figure 2** | Overview of the SIC classification in USTS and myxofibrosarcoma

**Supplementary figure 3** | Overview of immune contextures in USTS and myxofibrosarcoma

**Supplementary figure 4** | Association between immune infiltration and survival in USTS and myxofibrosarcoma

**Supplementary figure 5** | The effect of radiotherapy on USTS and myxofibrosarcoma

**Supplementary table 1** | Imaging mass cytometry marker panel

**Supplementary table 2** | Cell types and used lineage markers for the imaging mass cytometry analysis

**Supplementary table 3** | Univariate cox proportional hazard results for disease-specific and metastasis-free survival in USTS

**Supplementary table 4** | Univariate and multivariate cox proportional hazard results for disease-specific and metastasis-free survival in myxofibrosarcomas

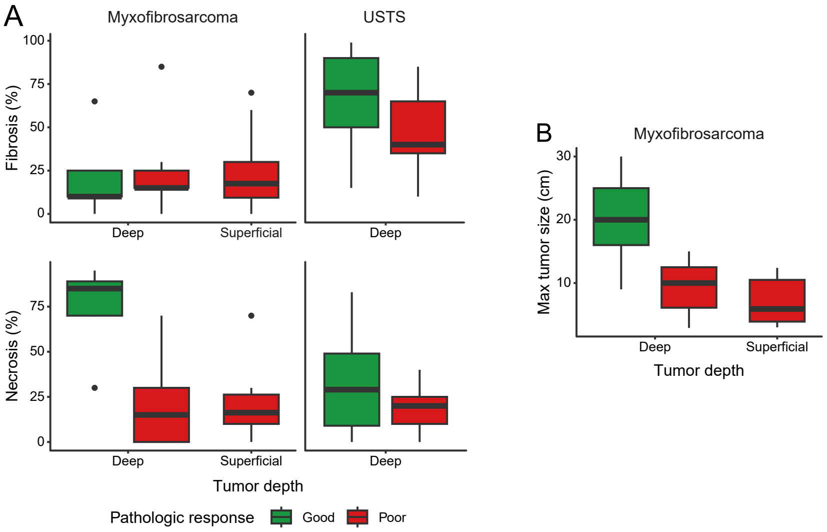

A
Myxofibrosarcoma
USTS
Fibrosis (%)
100
75
50
25
0
Deep
Superficial
Deep
Necrosis (%)
75
50
25
0
Deep
Superficial
Deep
Tumor depth
Pathologic response
Good
Poor
B
Myxofibrosarcoma
Max tumor size (cm)
30
20
10
Deep
Superficial
Tumor depth

**Supplementary Figure 1. Association between tumor depth, tumor size and necrosis in myxofibrosarcoma**. A) Boxplots displaying the association between pathologic response (<%5 vital tumor), fibrosis, necrosis, tumor depth and diagnosis. B) Boxplots presenting the association between max tumor size, pathologic response and tumor depth in myxofibrosarcoma. Abbreviations: USTS = undifferentiated soft tissue sarcoma.

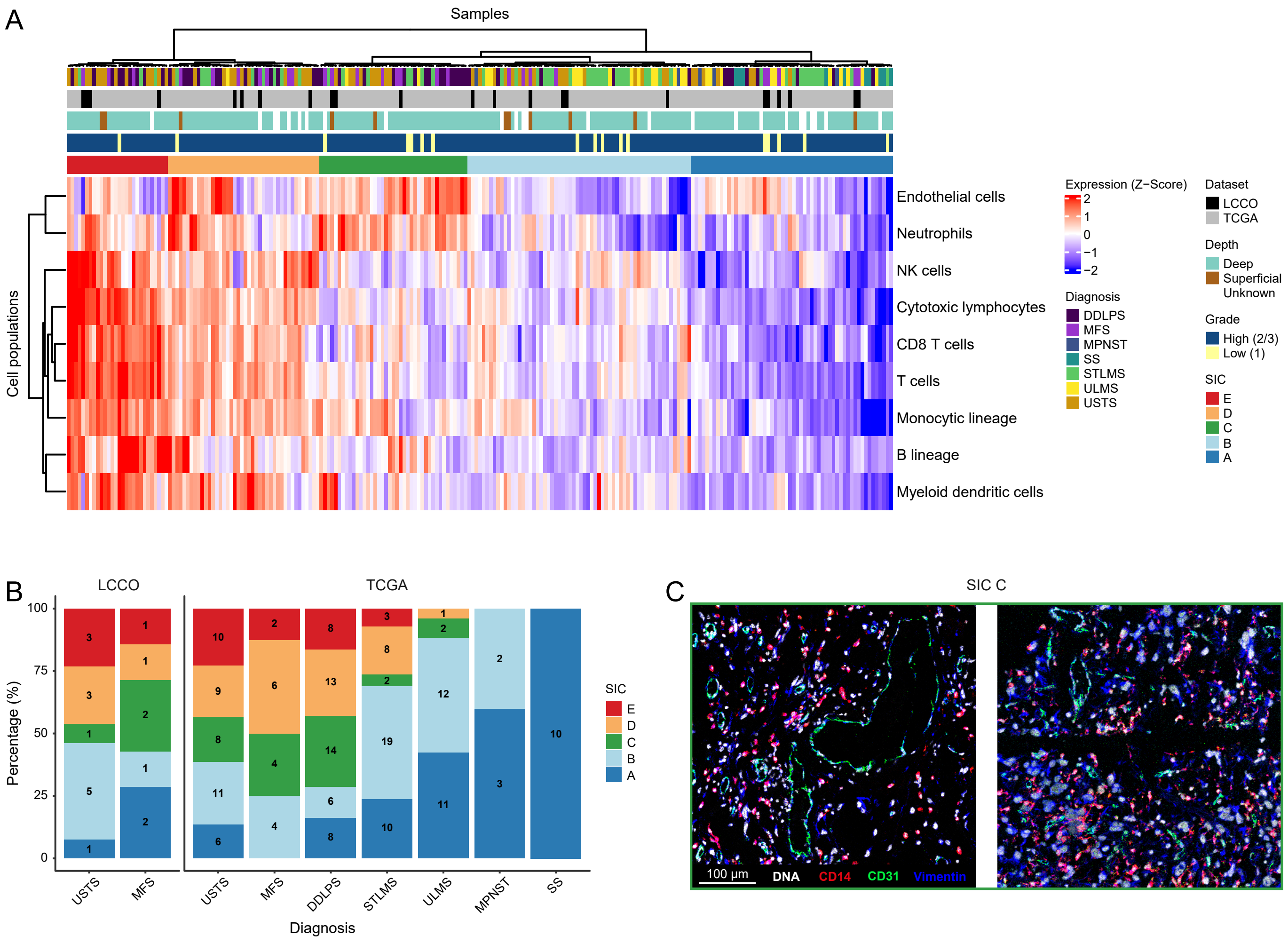
A
Samples
Cell populations
Endothelial cells
Neutrophils
NK cells
Cytotoxic lymphocytes
CD8 T cells
T cells
Monocytic lineage
B lineage
Myeloid dendritic cells
Expression (Z−Score)
2
1
0
−1
−2
Diagnosis
DDLPS
MFS
MPNST
SS
STLMS
ULMS
USTS
Dataset
LCCO
TCGA
Depth
Deep
Superficial
Unknown
Grade
High (2/3)
Low (1)
SIC
E
D
C
B
A
B
LCCO
TCGA
Percentage (%)
100
75
50
25
0
USTS
MFS
USTS
MFS
DDLPS
STLMS
ULMS
MPNST
SS
Diagnosis
C
SIC C
100 μm
DNA
CD14
CD31
Vimentin

**Supplementary figure 2. Overview of the SIC classification in USTS and myxofibrosarcoma.** A) Heatmap presenting the estimated cell population Z-score of the microenvironment cell population counter (MCP) in our samples and The Cancer Genome Atlas (TCGA) samples, resulting in the Sarcoma Immune Classes (SIC). Annotations include diagnosis, dataset, tumor depth, tumor grade and SIC classification. B) Bar plots illustrating the percentage of the SIC classification for each diagnosis across dataset, excluding low-grade tumors. The number of samples in each cluster is indicated within the bars. C) Representative imaging mass cytometry images of two SIC C (highly vascularized) myxofibrosarcomas. Abbreviations: DDLPS = dedifferentiated liposarcoma; MFS = myxofibrosarcoma; MPNST = malignant peripheral nerve sheath tumor; SS = synovial sarcoma; STLMS = soft tissue leiomyosarcoma; ULMS = uterine leiomyosarcoma; LCCO = Leiden Center for Computational Oncology.

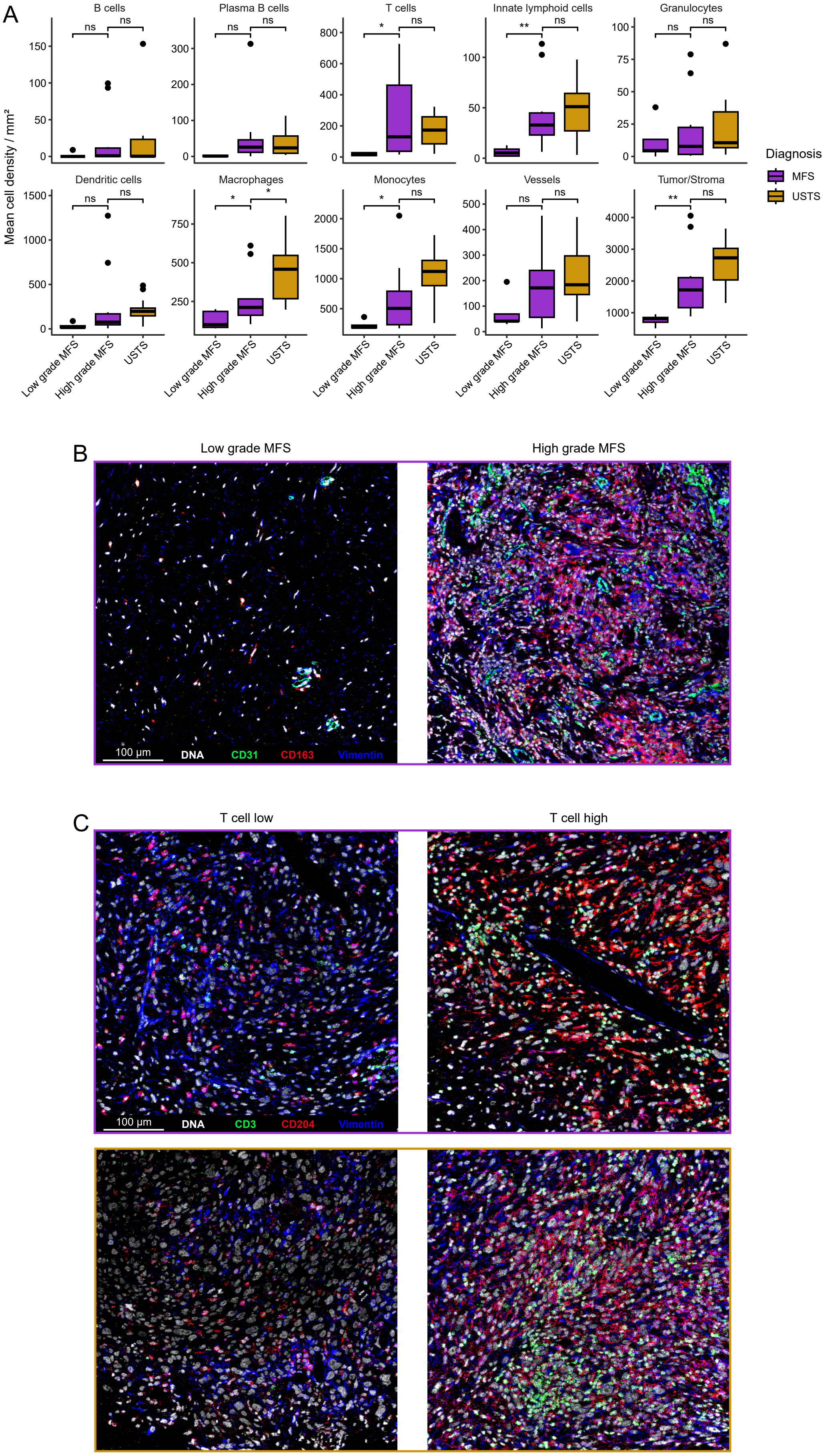

A
B cells
Plasma B cells
T cells
Innate lymphoid cells
Granulocytes
Dendritic cells
Macrophages
Monocytes
Vessels
Tumor/Stroma
Mean cell density / mm²
Diagnosis
MFS
USTS
Low grade MFS
High grade MFS
USTS
B
Low grade MFS
High grade MFS
100 µm
DNA
CD31
CD163
Vimentin
C
T cell low
T cell high
100 µm
DNA
CD3
CD204
Vimentin

**Supplementary figure 3. Overview of immune contextures in USTS and myxofibrosarcoma.** A) Boxplots presenting the mean cell density of major cell types for low-grade myxofibrosarcoma, high-grade myxofibrosarcoma and undifferentiated soft tissue sarcomas (USTS). Differences between low-grade and high-grade myxofibrosarcoma, as well as high-grade myxofibrosarcoma and USTS were evaluated with a student's t-test. ns = not significant, * = $P < 0.05$, ** = $P < 0.01$. B) Example imaging mass cytometry images of a low-grade and a high-grade myxofibrosarcoma, highlighting the difference in cellularity. The image displays tumor/stromal cells in blue (vimentin), vessels in green (CD31) and myeloid cells in red (CD163). C) Example imaging mass cytometry images of a T cell low and a T cell high, high-grade myxofibrosarcoma and USTS. The images display T cells in green (CD3), tumor/stromal cells in blue (vimentin) and myeloid cells in red (CD204). Abbreviations: MFS = myxofibrosarcoma.

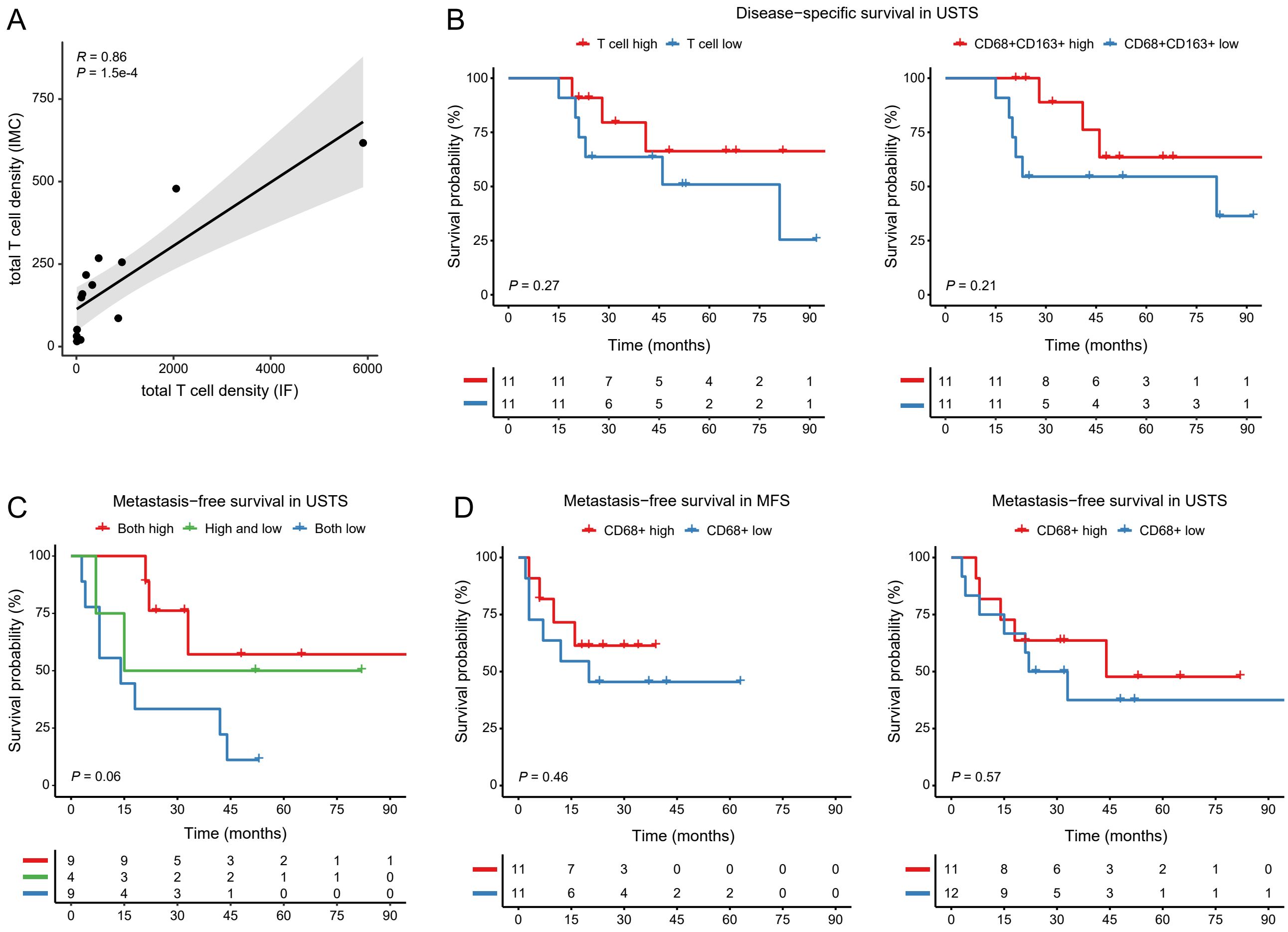
A
R = 0.86
P = 1.5e-4
total T cell density (IMC)
total T cell density (IF)
B
Disease−specific survival in USTS
T cell high
T cell low
CD68+CD163+ high
CD68+CD163+ low
Survival probability (%)
P = 0.27
P = 0.21
Time (months)
C
Metastasis−free survival in USTS
Both high
High and low
Both low
P = 0.06
D
Metastasis−free survival in MFS
Metastasis−free survival in USTS
CD68+ high
CD68+ low
P = 0.46
P = 0.57

**Supplementary figure 4. Association between immune infiltration and survival in USTS and myxofibrosarcoma.** A) Correlation plot presenting the positive correlation between T cell densities as detected with imaging mass cytometry (IMC) compared to immunofluorescence (IF). B) Survival analysis of the disease-specific survival in undifferentiated soft tissue sarcoma (USTS), based on the T cell and CD68+CD163+ macrophage groups. C) Survival analysis of the metastasis-free survival in USTS, grouped based on the combined T cell and CD68+CD163+ macrophage infiltration. D) Survival analysis of the metastasis-free survival in both myxofibrosarcoma and USTS, grouped based on the CD68+ macrophage infiltration. The significance of the Kaplan-Meier curves is presented by the log-rank *P* values. Abbreviations: MFS = myxofibrosarcoma.

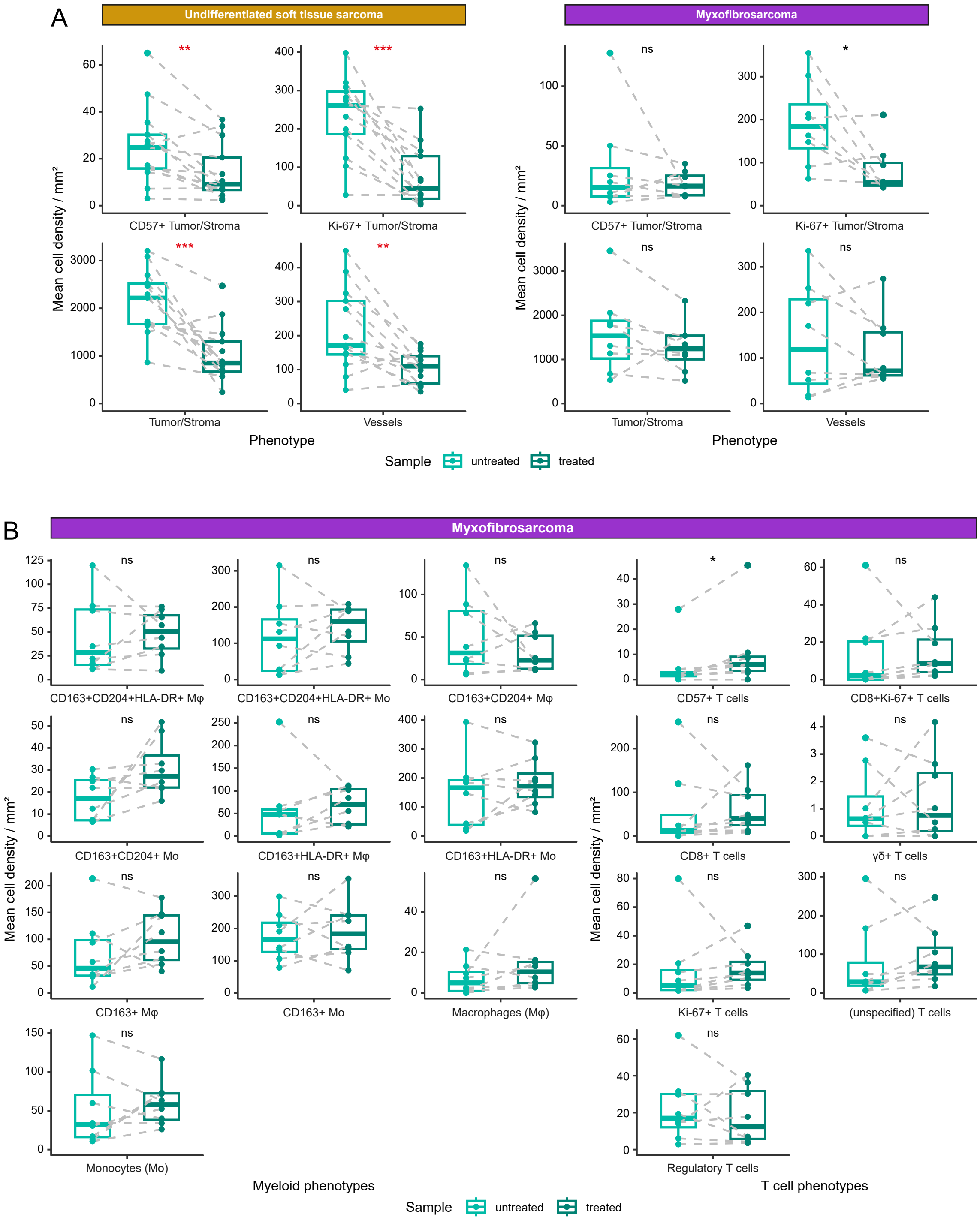
A
Undifferentiated soft tissue sarcoma
Myxofibrosarcoma
Mean cell density / mm²
CD57+ Tumor/Stroma
Ki-67+ Tumor/Stroma
Tumor/Stroma
Vessels
**
***
***
**
ns
*
ns
ns
Phenotype
Sample
untreated
treated
B
Myxofibrosarcoma
CD163+CD204+HLA-DR+ Mφ
CD163+CD204+HLA-DR+ Mo
CD163+CD204+ Mφ
CD57+ T cells
CD8+Ki-67+ T cells
CD163+CD204+ Mo
CD163+HLA-DR+ Mφ
CD163+HLA-DR+ Mo
CD8+ T cells
γδ+ T cells
CD163+ Mφ
CD163+ Mo
Macrophages (Mφ)
Ki-67+ T cells
(unspecified) T cells
Monocytes (Mo)
Regulatory T cells
Myeloid phenotypes
T cell phenotypes
Sample
untreated
treated

**Supplementary figure 5. The effect of radiotherapy on USTS and myxofibrosarcoma**. A) Paired boxplots presenting the other statistically significant alterations to the USTS immune microenvironment after radiotherapy. The same phenotypes are shown for myxofibrosarcoma as a means of comparison. The significance level was evaluated with a student's t-test followed by a Benjamini-Hochberg false discovery rate (FDR) correction. The significance is indicated per phenotype and FDR-significant phenotypes are indicated in red. ns = not significant, * = $P < 0.05$, ** = $P < 0.01$, *** = $P < 0.001$. B) Comparison of pre- vs post-treatment immune cell phenotypes in myxofibrosarcomas, presented in paired boxplots. The same phenotypes are presented in **Figure 4** for USTS. The significance is indicated per phenotype and FDR-significant phenotypes are indicated in red. Abbreviations: MFS = myxofibrosarcoma; Mϕ = macrophages; Mo = monocytes; HR = hazard ratio; CI = confidence interval.

**Supplementary table 1. Imaging mass cytometry marker panel.** Ab = antibody, ON = overnight, PDPN = podoplanin.

| | Target | Clone | Metal | Incubation Time | Temp | Dilution (x) |
|---|---|---|---|---|---|---|
| Lymphoid | CD103 | EPR4166(2) | 168 Er | 5h | RT | 50 |
| | CD19 | D4V4B | 172 Yb | 5h | RT | 100 |
| | CD20 | H1 | 142 Nd | Overnight | 4C | 100 |
| | CD27 | EPR8569 | 175 Lu | Overnight | 4C | 50 |
| | CD3 | EP449E | 153 Eu | Overnight | 4C | 50 |
| | CD38 | EPR4106 | 169 Tm | Overnight | 4C | 100 |
| | CD4 + 2nd AB | EPR6855 | 145 Nd | Indirect ON | 4C | 100 |
| | CD7 | EPR4242 | 174 Yb | 5h | RT | 100 |
| | CD8a | D8A8Y | 146 Nd | 5h | RT | 50 |
| | FOXP3 | D608R | 159 Tb | Overnight | 4C | 50 |
| | TCRγδ + 2nd Ab | H41 | 148 Nd | Indirect ON | 4C | 50 |
| Myeloid | CD11b | D6X1N | 144 Nd | 5h | RT | 100 |
| | CD11c | EP1347Y | 176 Yb | 5h | RT | 100 |
| | CD14 | D7A2T | 163 Dy | 5h | RT | 100 |
| | CD15 | MC480 | 171 Yb | Overnight | 4C | 100 |
| | CD163 | D6U1J | 173 Yb | 5h | RT | 50 |
| | CD204 | J5HTR3 | 164 Dy | 5h | RT | 50 |
| | CD68 | D4B9C | 143 Nd | Overnight | 4C | 100 |
| | HLA-DR | TAL 1B5 | 141 Pr | 5h | RT | 100 |
| Tumor/Stroma | CD31 | 89C2 | 147 Sm | Overnight | 4C | 100 |
| | CD39 | EPR20627 | 157 Gd | 5h | RT | 100 |
| | CD45 | D9M8I | 149 Sm | Overnight | 4C | 50 |
| | CD45RO | UCHL1 | 165 Ho | Overnight | 4C | 100 |
| | CD56 | E7X9M | 167 Er | 5h | RT | 100 |
| | CD57 | HNK-1 / Leu-7 | 151 Eu | Overnight | 4C | 100 |
| | D2-40 (PDPN) | D2-40 | 166 Er | Overnight | 4C | 100 |
| | Keratin | C11 and AE1/AE3 | 198 Pt | Overnight | 4C | 50 |
| | TGF-β | TB21 | 115 In | 5h | RT | 100 |
| | Vimentin | D21H3 | 194 Pt | Overnight | 4C | 50 |
| | β-Catenin | D10A8 | 89 Y | Overnight | 4C | 100 |
| Activation | Granzyme B | D6E9W | 150 Nd | 5h | RT | 100 |
| | ICOS | D1K2T(TM) | 161 Dy | 5h | RT | 50 |
| | IDO | D5J4E(TM) | 162 Dy | Overnight | 4C | 100 |
| | Ki-67 | 8D5 | 152 Sm | Overnight | 4C | 100 |
| | LAG-3 | D2G40(TM) | 155 Gd | 5h | RT | 50 |
| | PD-1 | D4W2J | 160 Gd | 5h | RT | 50 |
| | PD-L1 | E1L3N(R) | 156 Gd | Overnight | 4C | 50 |
| | Tbet | 4B10 | 170 Er | 5h | RT | 50 |
| | TIM-3 | D5D5R(TM) | 154 Sm | 5h | RT | 100 |
| | VISTA | D1L2G(TM) | 158 Gd | 5h | RT | 100 |
| DNA | Histone H3 | D1H2 | 209 Bi | Overnight | 4C | 50 |

**Supplementary table 2. Cell types and used lineage markers for the imaging mass cytometry analysis.**

| Phenotype | Cell type | Lineage markers |
|---|---|---|
| B cells | B cells | CD20+ |
| Plasma B cells | Plasma B cells | CD38+ |
| HLA-DR+<br>CD11c+HLA-DR+ | Dendritic cells | CD14-CD68- |
| Granulocytes | Granulocytes | CD15+ |
| CD56+<br>CD57+<br>Innate lymphoid cells<br>Ki-67+ | Innate lymphoid cells | CD3-CD7+ |
| CD163+CD204+<br>CD163+HLA-DR+<br>CD163+<br>CD163+CD204+HLA-DR+<br>Macrophages | Macrophages | CD68+ |
| CD163+CD204+HLA-DR+<br>CD163+CD204+<br>CD163+HLA-DR+<br>CD163+<br>Monocytes | Monocytes | CD14+CD68- |
| CD57+<br>CD8+Ki-67+<br>CD8+<br>γδ+<br>Ki-67+<br>T cells<br>Regulatory T cells | T cells | CD3+ |
| CD56+<br>CD57+<br>PDPN+<br>Ki-67+<br>Tumor/Stroma | Tumor/Stroma | Vimentin+lineage markers- |
| Vessels | Vessels | CD31+ |

**Supplementary table 3. Univariate cox proportional hazard results for disease-specific and metastasis-free survival in USTS.** The survival analysis includes 30 undifferentiated soft tissue sarcoma (USTS) patients who were neoadjuvantly treated with radiotherapy. Surgical margin was excluded because all USTS patients had R0 resection margins. Pathologic response was considered <5% vital tumor after treatment. Abbreviations: CI = confidence interval; HR = hazard ratio.

| USTS | | |
|---|---|---|
| **Disease-specific survival** | **Univariate** | |
| Variable | **HR (95% CI)** | **log-rank *P*** |
| **Age at diagnosis** | 1.01 (0.98 - 1.06) | 0.7 |
| **Sex (Female)** | | |
| Male | 0.62 (0.19 - 2.03) | 0.4 |
| **Location (Lower extremities)** | | |
| Other[a] | 1.3e-08 (0 - Inf) | 0.3 |
| Upper extremities | 2.1 (0.43 - 10) | |
| **Max tumor size (cm)[b]** | 1.05 (0.96 - 1.2) | 0.3 |
| **Pathologic response (Good)** | | |
| Poor | 0.46 (0.14 - 1.5) | 0.2 |

a: Other includes tumors from the trunk and the head & neck area.
B: Tumor size was missing for one tumor from the head & neck area.

| USTS | | |
|---|---|---|
| **Metastasis-free survival** | **Univariate** | |
| Variable | **HR (95% CI)** | **log-rank *P*** |
| **Age at diagnosis** | 1.02 (0.98 - 1.06) | 0.3 |
| **Sex (Female)** | | |
| Male | 0.62 (0.24 - 1.6) | 0.3 |
| **Location (Lower extremities)** | | |
| Other | 1.3e-08 (0 - Inf) | 0.06 |
| Upper extremities | 2.8 (0.88 - 8.7) | |
| **Max tumor size (cm)** | 1.03 (0.95 - 1.1) | 0.4 |
| **Pathologic response (Good)** | | |
| Poor | 0.79 (0.31 - 2) | 0.6 |

**Supplementary table 4. Univariate and multivariate cox proportional hazard results for disease-specific and metastasis-free survival in myxofibrosarcomas.** The survival analysis includes 24 high-grade myxofibrosarcoma patients who were neoadjuvantly treated with radiotherapy. Pathologic response was considered <5% vital tumor after treatment. Abbreviations: CI = confidence interval; HR = hazard ratio.

| Myxofibrosarcoma | | | | |
|---|---|---|---|---|
| **Disease-specific survival** | **Univariate** | | **Multivariate** | |
| Variable | **HR (95% CI)** | **log-rank *P*** | **HR (95% CI)** | **log-rank *P*** |
| **Age at diagnosis** | 1.1 (0.98 - 1.3) | 0.1 | | |
| **Sex (Female)** | | | | |
| Male | 1.4 (0.26 - 7.7) | 0.7 | | |
| **Margin (R0)** | | | | |
| R1/R2 | 1.3 (0.15 - 12) | 0.8 | | |
| **Location (Lower extremities)** | | | | |
| Trunk | 0.58 (0.066 - 5.1) | 0.4 | | |
| Upper extremities | 3.3 (0 - Inf) | | | |
| **Tumor depth (Deep)** | | | | |
| Superficial | 7.1e-10 (0 - Inf) | **0.03 *** | 2.4e-09 (0 - Inf) | |
| **Max tumor size** | **1.3 (1.1 - 1.5) **** | **4e-04 ***** | 1.2 (0.97 - 1.4) | **0.002 **** |
| **Pathologic response (Good)** | | | | |
| Poor | **0.096 (0.015 - 0.6) *** | **0.002 **** | 0.57 (0.054 - 6) | |

| Myxofibrosarcoma | | | | |
|---|---|---|---|---|
| **Metastasis-free survival** | **Univariate** | | **Multivariate** | |
| Variable | **HR (95% CI)** | **log-rank *P*** | **HR (95% CI)** | **log-rank *P*** |
| **Age at diagnosis** | 1 (0.96 - 1.1) | 0.4 | | |
| **Sex (Female)** | | | | |
| Male | 1.8 (0.45 - 6.9) | 0.4 | | |
| **Location (Lower extremities)** | | | | |
| Trunk | 0.45 (0.56 - 3.6) | 0.6 | | |
| Upper extremities | 0.45 (0.56 - 3.6) | | | |
| **Margin (R0)** | | | | |
| R1/R2 | 0.79 (0.1 - 6.3) | 0.8 | | |
| **Tumor depth (Deep)** | | | | |
| Superficial | 0.14 (0.17 - 1.08) | **0.03 *** | 0.22 (0.026 - 1.9) | |
| **Max tumor size (cm)** | **1.2 (1.04 - 1.3) **** | **0.004 **** | 1.1 (0.97 - 1.2) | **0.005 **** |
| **Pathologic response (Good)** | | | | |
| Poor | **0.17 (0.044 - 0.66) *** | **0.004 **** | 0.46 (0.087 - 2.4) | |
